# SParticle, an algorithm for the analysis of filamentous microorganisms in submerged cultures

**DOI:** 10.1101/159475

**Authors:** Joost Willemse, Ferhat Büke, Dino van Dissel, Sanne Grevink, Dennis Claessen, Gilles P. van Wezel

## Abstract

Streptomycetes are filamentous bacteria that produce a plethora of bioactive natural products and industrial enzymes. Their mycelial lifestyle typically results in high heterogeneity in bioreactors, with morphologies ranging from fragments and open mycelial mats to dense pellets. There is a strong correlation between morphology and production in submerged cultures, with small and open mycelia favoring enzyme production, while most antibiotics are produced mainly in pellets. Here we describe SParticle, a *Streptomyces* Particle analysis method that combines whole slide imaging with automated image analysis to characterize the morphology of submerged grown *Streptomyces* cultures. SParticle allows the analysis of over a thousand particles per hour, offering a high throughput method for the imaging and statistical analysis of mycelial morphologies. The software is available as a plugin for the open source software ImageJ and allows users to create custom filters for other microbes. Therefore, SParticle is a widely applicable tool for the analysis of filamentous microorganisms in submerged cultures.

## INTRODUCTION

Streptomycetes are Gram-positive bacteria that have a complex mycelial lifestyle, whereby they reproduce via sporulation (Claessen 2014; Chater and Losick 1997). Streptomycetes and other members of the phylum of actinomycetes are of great medical, biotechnological and agricultural importance due to their ability to produce a wide variety of bioactive natural products (Barka 2016; Bérdy 2005; Hopwood 2007). Furthermore, their saphrophytic lifestyle allows them to degrade almost all known, natural bio-polymers (Omura 1992). Morphology and growth characteristics may vary strongly, based on strain-specific traits that are yet poorly understood (Jakimowicz and van Wezel 2012). During growth in submerged cultures, *Streptomyces* species form mycelial structures, which can be classified in three groups based on the different phenotypes (van Dissel 2014). Many streptomycetes form extended mycelial networks, which often self-aggregate into dense cellular structures called pellets (Celler 2012; Meyerhoff 1995). Other morphologies include mycelial mats (caused by open growth) and small hyphal fragments (Tresner 1967). *Streptomyces* can be further subdivided into those that sporulate in liquid cultures and those that do not (Girard 2013; Glazebrook 1990; Kendrick and Ensign 1983). When grown under different conditions, growth rate and morphology are not just strain specific, but also change depending on the composition of the growth medium, pH, temperature, shear rate, dissolved oxygen concentration and inoculum (Tough and Prosser 1996; Cui 1998). The relationship between growth and morphology, on the one hand, and biomass accumulation and productivity on the other, is complicated, and optimal morphology varies from product to product (van Wezel 2009). While pellets are primarily physiologically active around the edges and therefore associated with slow growth, they promote antibiotic production, presumably correlated to nutrient limitation in the core of the pellets (van Dissel 2014; Wardell 2002). Conversely, smaller pellets and open mycelia are generally preferred morphologies for enzyme production, which may be explained by the fact that protein production preferentially takes place at apical sites (van Wezel 2006; Willemse 2012).

Detailed assessment of the morphological characteristics of a culture is important for rational strain design (Celler 2012). Attempts in the 1990s aimed at qualitatively assessing the morphology of filamentous microorganisms, applied a combination of microscopy with the advances in computational power for (semi-)automated image analysis to measure the size and other characteristics such as shape, density and complexity of pellets (Reichl 1992; Reichl 1990; Treskatis 1997; Pinto 2004; Barry and Williams 2011). A limitation of microscopy-based techniques is the trade-off between the throughput and image quality. Sufficient resolution is required to allow reliable image analysis, but higher magnification limits the number of particles that can be captured (Barry and Williams 2011; Papagianni 2014; Hardy 2017). Various methods have been tried to increase the sample size in image analysis methods (Cox and Thomas 1992; Packer and Thomas 1990). Flow cytometry using a Complex Object Parametric Analyzer and Sorter (COPAS) efficiently analyzes pellets by size and fluorescence, allowing up to several 100s of measurements per second (Petrus 2014). There are however technical limitations; firstly, detection of small mycelial fragments is unreliable because their extinction coefficient is close to the background value, and the same is true for hyphae at the edge of open mycelial aggregates, making the relation between time of flight and pellet size hard to calibrate. Secondly, the time of flight fails to take the shape of pellets into account. Since pellets are typically non-circular, COPAS measurements cannot be used for pellet-volume estimation. This is also reflected in the larger standard deviation often observed in cytometry measurements (Rønnest 2012). Thus, microscopy allows acquiring information about pellet shape, density and fragmentation, while flow cytometry is ideal for studying population traits, due to the high number of measurements.

To overcome the low sample size in microscopic measurements we applied whole slide imaging (WSI) combined with automated image analysis to increase the throughput without compromising image quality and thus reliability. WSI has been used by pathologists for more than a decade (Webster and Dunstan 2014). Imaging of a whole pathological sample and subsequent automated image analysis is becoming common practice in areas such as cancer research (Cooper 2015) and developments on staining techniques continuously expand the applicability of WSI (Gray 2015). So far, WSI has found limited application in microbial population measurements (Posch 2012), and no software is currently freely available. Here we describe a WSI-based method for automated imaging of *Streptomyces* mycelia. By capturing a high-resolution picture of a large area, our approach visualizes a large number of particles, and accurately identifies particles ranging from small mycelial fragments to large pellets. As our method is applicable to other filamentous micro-organisms, we expect that this free software will allow labs to improve the quantitatively and qualitatively description of microbial morphologies.

## MATERIALS AND METHODS

### Strains and culturing conditions

*Streptomyces coelicolor* FM145 (Willemse and van Wezel 2009) was used as reference strain in all experiments. FM145 is a derivative of *S. coelicolor* M145 (Bentley 2002) with reduced autofluorescence. The filamentous fungus *Aspergillus niger* N402, as well as the cyanobacteria *Gloeothece sp.* PCC 6909, *Chroococcidiopsis sp.* PCC 6712 and *Synechococcus elongatus* PCC 7942 were obtained from our in-house strain collection. *A. niger* N402 was grown at 30°C in complete medium, consisting of minimal medium supplemented with 1% yeast extract and 0.5% casamino acids (Vinck 2005). *Gloeothece sp.* PCC 6909, *Chroococcidiopsis sp.* PCC 6712 and *Synechococcus elongatus* PCC 7942 were cultured in BG-11 media as described (Voshol 2015). *Streptomyces* spores were collected from MS agar plates (Kieser 2000). Cfus (colony forming units) were determined by plating serial dilutions on MS agar plates. For submerged growth and pellet size measurement experiments, *S. coelicolor* was cultivated in spring-loaded 125 ml Erlenmeyer flasks containing 25 ml Tryptic Soy Broth media supplemented with 10.3% sucrose (TSBS). Cultures of *S. coelicolor* FM145 were inoculated with 10^6^ Cfu spores/ml unless stated otherwise. A one inch orbital shaker (New Brunswick) was used at 30°C with shaking at 200 rpm.

### Microscopy and software

For imaging of the samples, we used a setup consisting of a Zeiss inverted Axio Observer with automated XY-stage using a 10x objective and a Hamamatsu EM-CCD c9100-02 camera. Samples were taken from submerged cultures using a wide-gauge 200 µl pipette tip to allow big pellets to pass, and transferred to a microscope slide, covered with a 2×2cm square cover slip and sealed by using fast drying nail polish to prevent evaporation induced artefacts during imaging. The automated stage served to capture a multi-image phase-contrast mosaic, imaged by meandering from top left to bottom-right (Fig S1). The outer 2 mm of the cover slip were excluded from measurements to exclude an effect of the sealing material. By using a 20 by 20 grid, with 10% overlap of images, we were able to capture the ∼1.4×1.4 cm inner area of the slide, which takes ∼8 minutes to complete. The software was developed as a plugin for the ImageJ package (https://imagej.nih.gov/ij/index.html), which is freely available from the Leiden University ImageJ update site (https://imagej.net/List_of_update_sites).

## RESULTS

### Automated image analysis steps

We sought to develop software that allows researchers to reliably assess morphological parameters of submerged cultures of filamentous microorganisms at medium to high throughput scale. A major aim was to combine automated image analysis with automated whole slide imaging so as to allow assessing the morphology of mycelia in submerged cultures for any given culture within fifteen minutes in a statistically relevant manner.

#### Mosaic image building and image normalization

Submerged cultures were set up by inoculating TSBS cultures with 10^6^ cfu and grown for 24 h at 30°C. Subsequently, mosaic images of the samples were obtained using an automated stage microscope. To quickly analyze a large number of mycelia, mosaic images were created using the Axiovision software and stored as separate TIFF files. These files were reconstructed into large mosaic images thereby eliminating partial pellets at the borders of individual images. Furthermore, the images were normalized to correct for light fluctuations from the light source. Using a 20×20 grid, with 10% overlap of images, an approximately 1.4×1.4 cm inner area of the slide was captured in approximately 8 minutes. 400 images per slide were taken and analyzed in ImageJ. As a rule of thumb, 12 Gb of RAM is sufficient to analyze ∼325 Megapixel images, which means that an entire microscope slide can be analyzed as a single image. Sampling density was chosen such that most pellets and fragments are spatially separated, simplifying analysis of the objects and allowing brightness adjustment of each mosaic image using the background intensity determined as grey values. This method also allows analysis of fluorescence images, as well as analysis of different mycelial morphologies (Zacchetti 2016)

#### Determining the Region of Interest (ROI)

Thresholding was applied to separate candidate objects from the background, creating a binary image. Pixels with brightness values between 50 and 90 were assigned as background, while the remaining pixels were assigned as possible pellet edges and small mycelial fragments. The detection of small fragments was facilitated by phase-contrast imaging, where changes in light permeability are emphasized resulting in illuminated edges. After thresholding, area filling was applied to generate solid objects out of the detected edges (Fig. 1). Objects larger than 6×10^5^ µm^2^ (corresponding to an object with a diameter larger than 900 µm) were ignored to remove artifacts such as air bubbles, while objects smaller than 200 pixels were filtered out to remove small artifacts like cell debris or precipitated salts.

**Figure 1.**
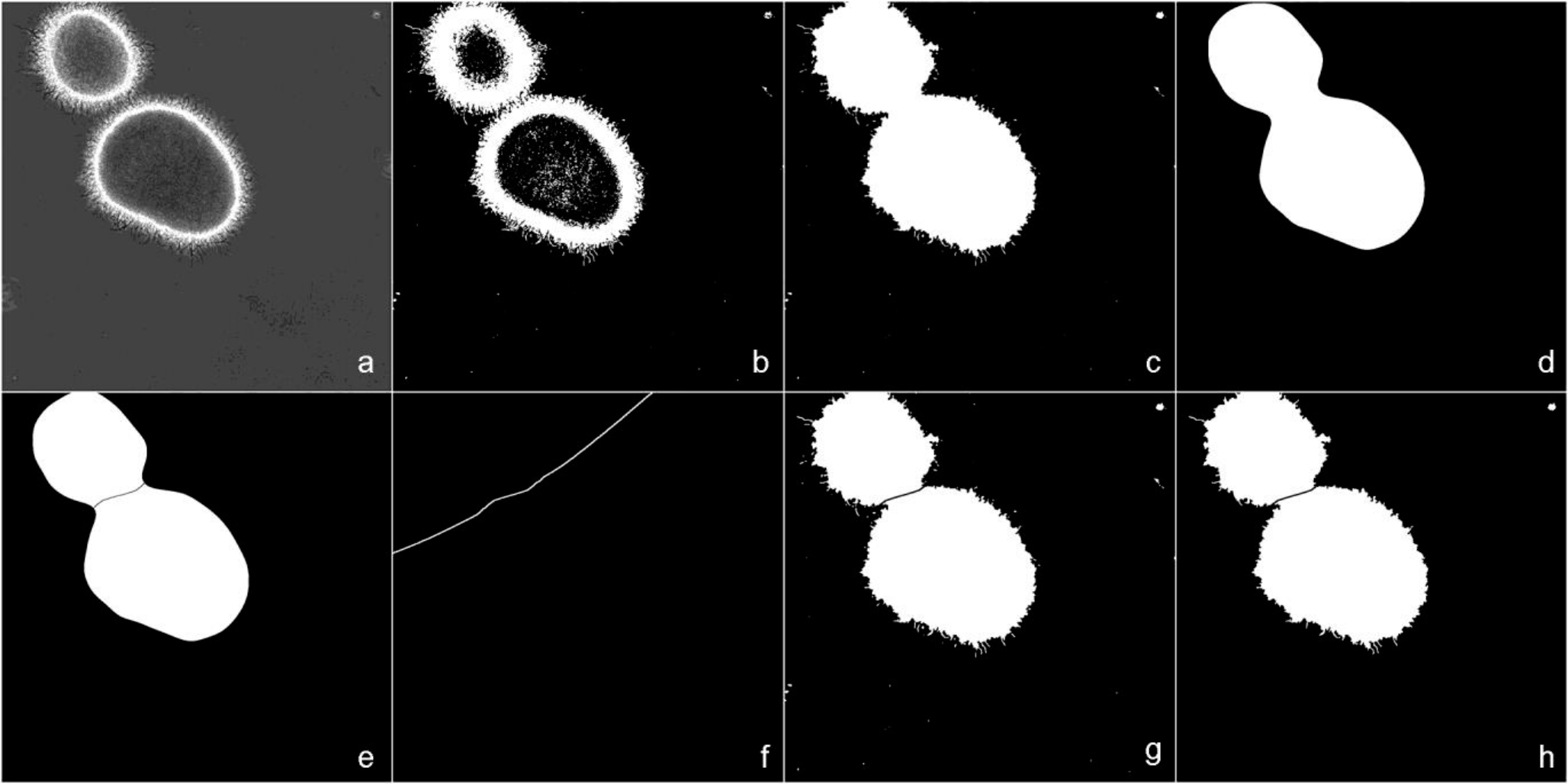
Steps of ROI Identification. The original image (a) is transformed into black & white (B&W b) using thresholding. Fully enclosed black areas are filled with white (c) and a Gaussian blur (d) is applied. Areas that are connected via “thin” bridges are separated via Watershedding (e) and a Voronoi transformation (f) is applied to find object areas. Identified area borders are subtracted from the filled B&W image (g) and small objects are deleted (h). Individual areas are then marked as ROIs, which are now ready for advanced filtering to further analyse the particles.

To maximize sample size and optimize the accuracy of the measurements, groups of particles were separated by water-shedding the mask after Gaussian blur. This approach separates adjacent objects as shown in Fig 1e. The risk of separating irregularly shaped pellets into two objects is minimized by only applying a separation if the interface between the two adjacent particles is smaller than 150 µm. Separation between adjacent objects is achieved by using a Voronoi transformation. This transformation divides the image into as many regions as there are objects in the total image (Fig. 1).

A rule set was applied on the ROIs to separate contaminants from pellets or mycelial fragments. For example, this filtering step eliminates large non-circular objects (e.g. cotton fibers) and objects deviating from the normal pellet density. For a complete set of rules, we refer to the Supplemental Methods. To allow a broader use of the plugin, a manually defined filter can be applied to any given set of ROIs. An additional macro tool is provided to create these manual filters. In this paper only the settings, optimized for streptomycetes, were used (see Supplemental Methods). Depending on the density of the sample, an image may contain up to 500 pellets and mycelial fragments. These are all identified and marked as separate ROIs and further measured. Saving all regions as well as the selected ones in separate files allows manual correction for false negatives if required.

#### Determining particle shape

After eliminating non-desirable objects from the potential list of particles, multiple parameters need to be measured to describe the objects in more detail. We chose the following parameters to describe the objects of interest: area, mean intensity, standard deviation of intensity, roundness, circularity and Feret’s diameter (an object’s diameter along its longest axis). In ImageJ, the area to be measured is rotated with 2 degree increments and Feret’s diameter is calculated vertically. The maximum measurement is taken as the longest distance within the object.

After processing by ImageJ, all selected ROIs were subjected to additional measurements. Initially the roughness was determined by dividing the perimeter of the object by the perimeter of the fitted ellipse. Furthermore, the morphological number, a dimensionless parameter which combines multiple morphological parameters and has been shown to correlate to productivity, was defined (Wucherpfennig 2011; Wucherpfennig 2013), and subsequently the fractal box mass dimension as well as the box surface dimension and the fractal quotient are determined (Obert 1990; Ryoo 1999). These three morphological pellet parameters distinguish different mycelial morphologies. Another circularity measurement based on Cartesian to Polar coordinate transformation (Stojmenovic 2013) was also determined. This method is more suited for estimating the circularity of a pellet, as other conventional methods use the perimeter length of an object. The object perimeter of a mycelial particle is highly dependent on outwardly protruding hyphae. Therefore, the number and length of the hyphae dictates conventional circularity calculations, making them unsuited to describe the circularity (see Fig. S2 for more details). We also implemented a band-intensity measurement, where donut shaped areas are placed that radiate from the center of the shape. The shapes of the donuts are corresponding to the shape of the ROI and each ROI is divided into 20 regions. By default, pellet images in phase contrast microscopy have relatively dark protruding hyphae, then a brighter ring at which the pellets dense region starts, followed by a darker interior. We make use of this by determining the peak intensity of the selected bands and then determining its positions relative to the edge of the pellet. This determines the width of the less dense outer perimeter of the pellet. The difference between the edge and the maximum allows determining the size of the low-density exterior in micrometers. Finally, we determine the outer intensity ratio to the inner intensity to determine how dense the pellets are, this parameter can be helpful in assisting to differentiate between pellet (with a dense/dark center) and mat formers (open center).

### From plugin to practice: imaging pellet formation by *Streptomyces coelicolor*

#### Optimizing the software for imaging of Streptomyces mycelia

Next, we optimized the microscope settings for the analysis of *Streptomyces* mycelia. A 10x magnification results in a pixel size of 780 nm, which suffices to distinguish mycelial fragments larger than 2 µm, which we chose as cut-off. We used this magnification throughout all experiments to minimize capture time while maintaining sufficient resolution. 42 WSI images were analyzed, resulting in the identification of more than 10,000 mycelial objects. To check the efficiency of the filter we manually confirmed or rejected all identified objects. This revealed 1.5 ± 0.46% false positives (average 262 correct objects per image detected, and 4 incorrect ones). Manual inspection of over 27,000 rejected objects revealed 1.06 ± 0.02% false negatives, in other words rejected objects that are proper mycelial objects (on average 658 correctly rejected objects per image detected, and 7 falsely rejected).

To test the reproducibility of the particles imaged by WSI combined with the ImageJ plugin SParticle, *S. coelicolor* was cultivated using the standard conditions described above. From a single flask three separate samples were analyzed to detect measurement variations. Comparing morphological descriptive parameters, including area, Feret diameter and circularity, no significant differences were found between the three replicate samples (Fig. 2a). This shows that the method reproducibly analyzed the particles. Importantly, adding an identical inoculum to cultures grown in independent shake flasks also resulted in highly similar morphologies (Fig. 2b).

**Figure 2.**
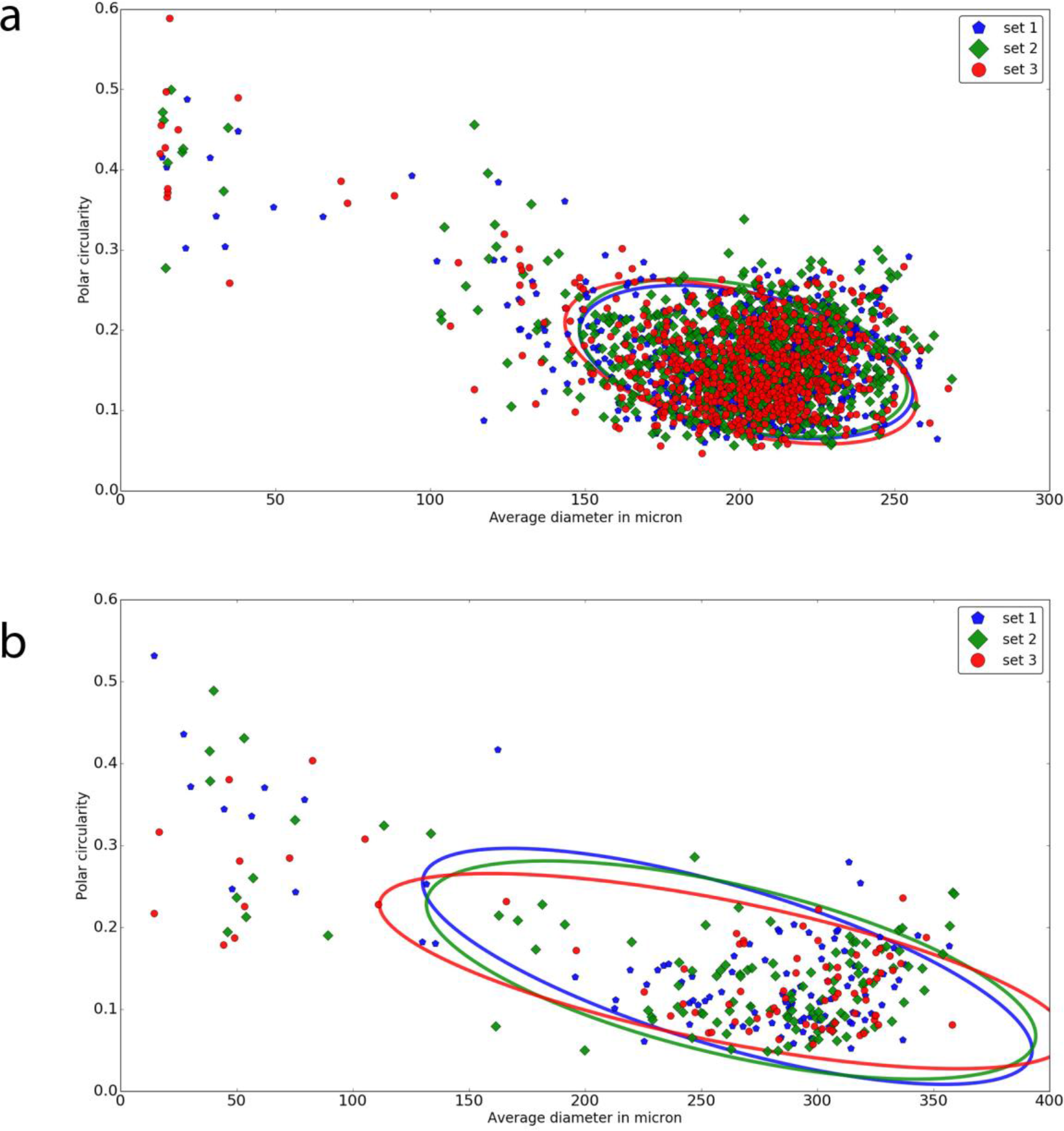
Reproducibility of measurements performed with SParticle. (a) Analysis of particles from a submerged culture, performed in triplicate, revealed no significant differences in the average diameter versus the polar circularity between replicates. The circles represent the 95% confidence intervals as determined with the error ellipse plotter (Kington 2012). (b) Likewise, no differences were observed when samples were taken after 24 h from different culture flasks.

#### Correlating inoculum and pellet size

To assess if we could distinguish between samples with a distinct pellet size distribution, cultures were inoculated at different spore densities. This was based on the observation that the inoculum density correlates to pellet size, where a higher number of spores results in a reduced average pellet size (Vecht-Lifshitz 1990). For this, 25 mL TSBS cultures were inoculated with 10^5^, 10^6^ or 10^7^ cfu of *S. coelicolor* spores and samples were taken after 24 h and 48 h of growth and analyzed by WSI. After 24 h, no statistically relevant difference was observed between the three inoculation densities, which all yielded pellets with a similar diameter vs. polar circularity (95% CI; Fig. 3a). After 48 h, small mycelial fragments represented a major fraction of all detected particles (Fig. 3b, left hand side). Fragmentation was observed at all inoculation densities, leading to the establishment of a population of small mycelial fragments and a separate population of larger particles. Notably, the average diameter of large particles was significantly different between the three cultures and depended on the starting inoculum (Fig. 3e), with T-test p-values of 1.1×10^−11^, 1.04×10^−74^, 1.46×10^−63^ for 10^5^ vs 10^6^, 10^5^ vs 10^7^ and 10^6^ vs 10^7^, respectively. These differences between inoculum size and average pellet size, measured in the population from which small fragments were excluded, are in agreement with literature (Vecht-Lifshitz 1990), which underlines the applicability of SParticle for the analysis of pellet morphologies.

**Figure 3.**
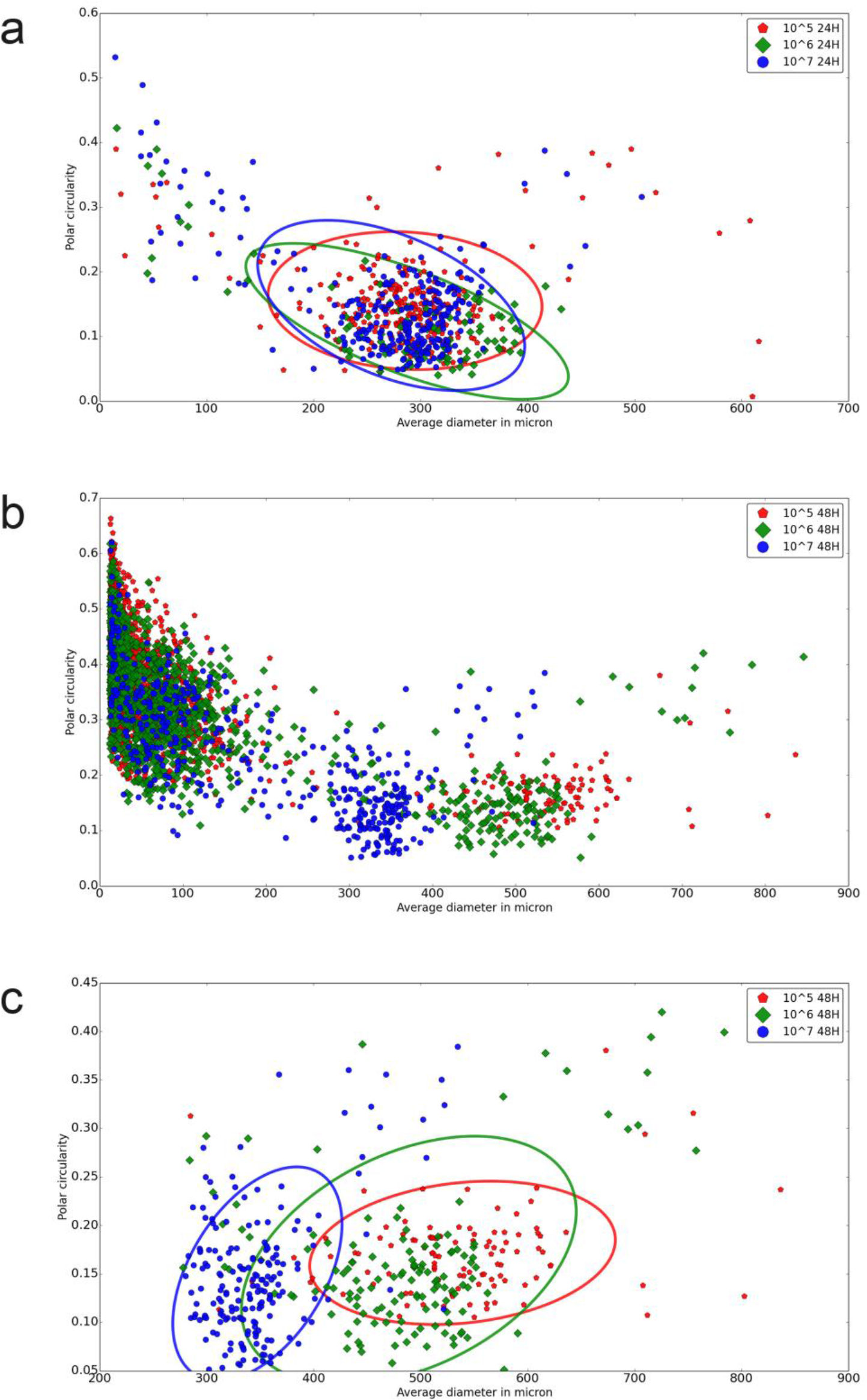
Correlation between inoculum and pellet morphology. (a) Inoculation density versus pellet size at 24 h. Measurement on different inoculation densities after 24 h reveals no differences in pellet morphology in early growth. For increasing inoculation density the amounts of pellets analyzed were 276, 105, and 292 respectively. Plotted are the average diameter versus the polar circularity, whereby the circles represent the 95% confidence intervals. (b) Inoculation density versus pellet size at 48 h. Measurements after 48 h indicate a large part of the population to be present as mycelial fragments and small pellets (<150 micron in length). The number of particles analyzed for increasing inoculation density were 1980, 1349, and 520 respectively. (c) Analysis of pellets from 48 h cultures confirms an inverse relationship between pellet size and inoculation density. The number of pellets analyzed for increasing inoculation density were 112, 149, and 213 respectively. Plotted are the average diameter versus the polar circularity; circles represent 95% confidence intervals.

#### Influence of shaker speed and orbital size on pellet size

Analysis of 24 h cultures at various shaker speeds and orbital sizes revealed no differences between baffled flasks or flasks with stainless steel spring coils (Fig. 4a). All other growth conditions were kept the same as described above. However, large differences were seen when flasks without baffles or springs were used (Fig. 4b). Decreasing the shaker speed from 200 rpm to 100 rpm led to a three-fold increase in average pellet diameter (Fig. 4c). We did not observe significant differences (p 0.09) between 125 ml flasks and 250 ml flasks filled with equal relative volumes (Fig. 4d), in line with previous observations (Mehmood 2011).

**Figure 4.**
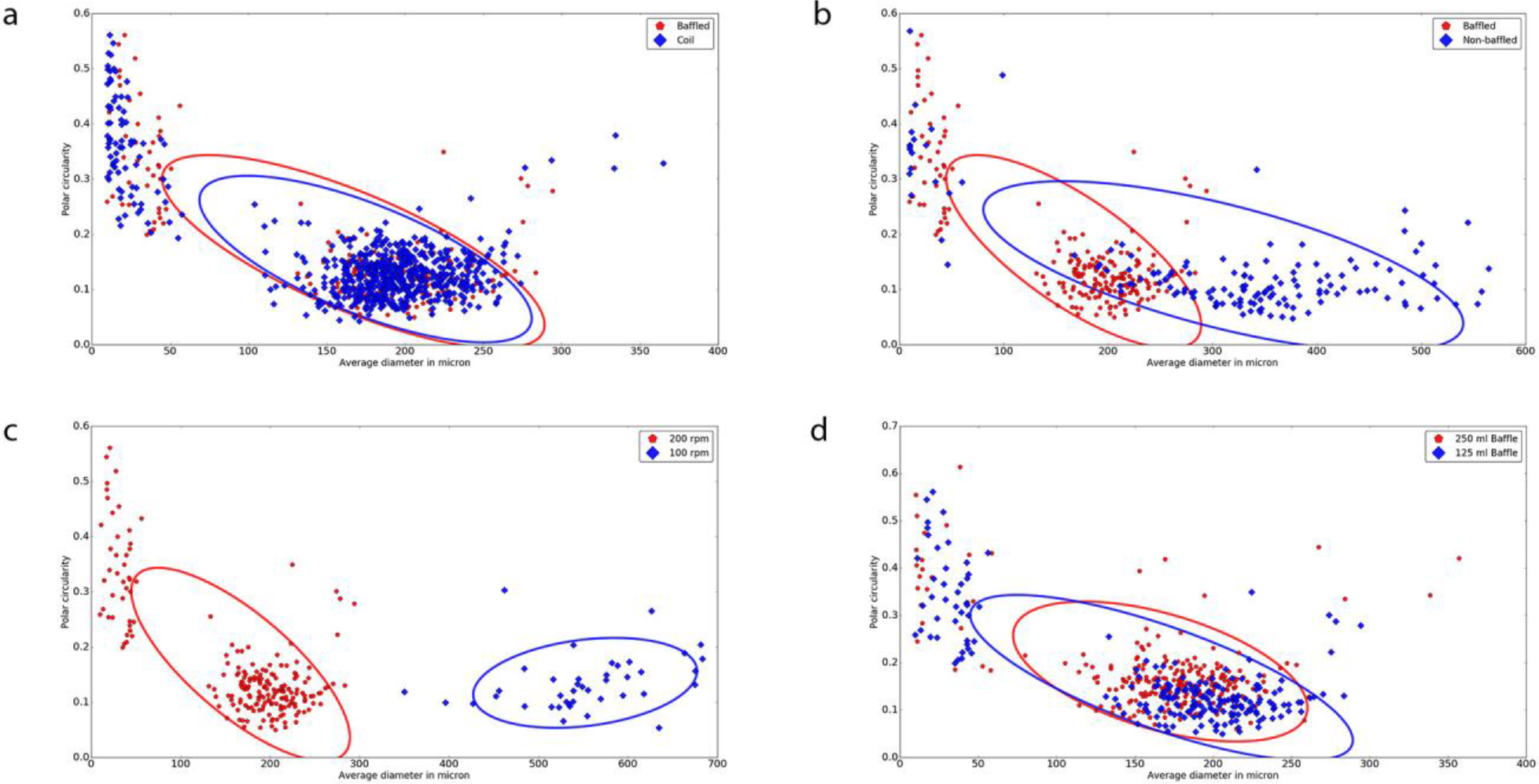
Shake speed and orbital size versus pellet size. Differences between baffles, coils, common flasks, shake speeds and flask volume. Based on these data we see no differences between coils and baffles (a) or between different inoculation volumes (d). However clear differences in pellet size were observed between unbaffled and baffled flasks (b) and when the orbital shaking speed was adjusted (c). Plotted are the average diameter versus the polar circularity; circles represent the 95% confidence intervals.

#### Application to other filamentous organisms

To establish if SParticle can also be used to analyze the morphology of other filamentous microorganisms we analyzed the mycelia of *Aspergillus niger* N402 grown for 24 h and *Gloeothece* sp. PCC 6909, *Chroococcidiopsis* sp. PCC 6712 and *Synechococcus elongatus* PCC 7942 grown for 120 h. From the image data, the software identified small and large pellets or assemblies of *A. niger,* Gloeothece sp. PCC 6909, *Chroococcidiopsis* sp. PCC 6712 and S. *elongatus* PCC 7942 (Fig. 5). This further underlines the applicability of SParticle for the analysis of the morphology of filamentous microorganisms in submerged cultures.

**Figure 5.**
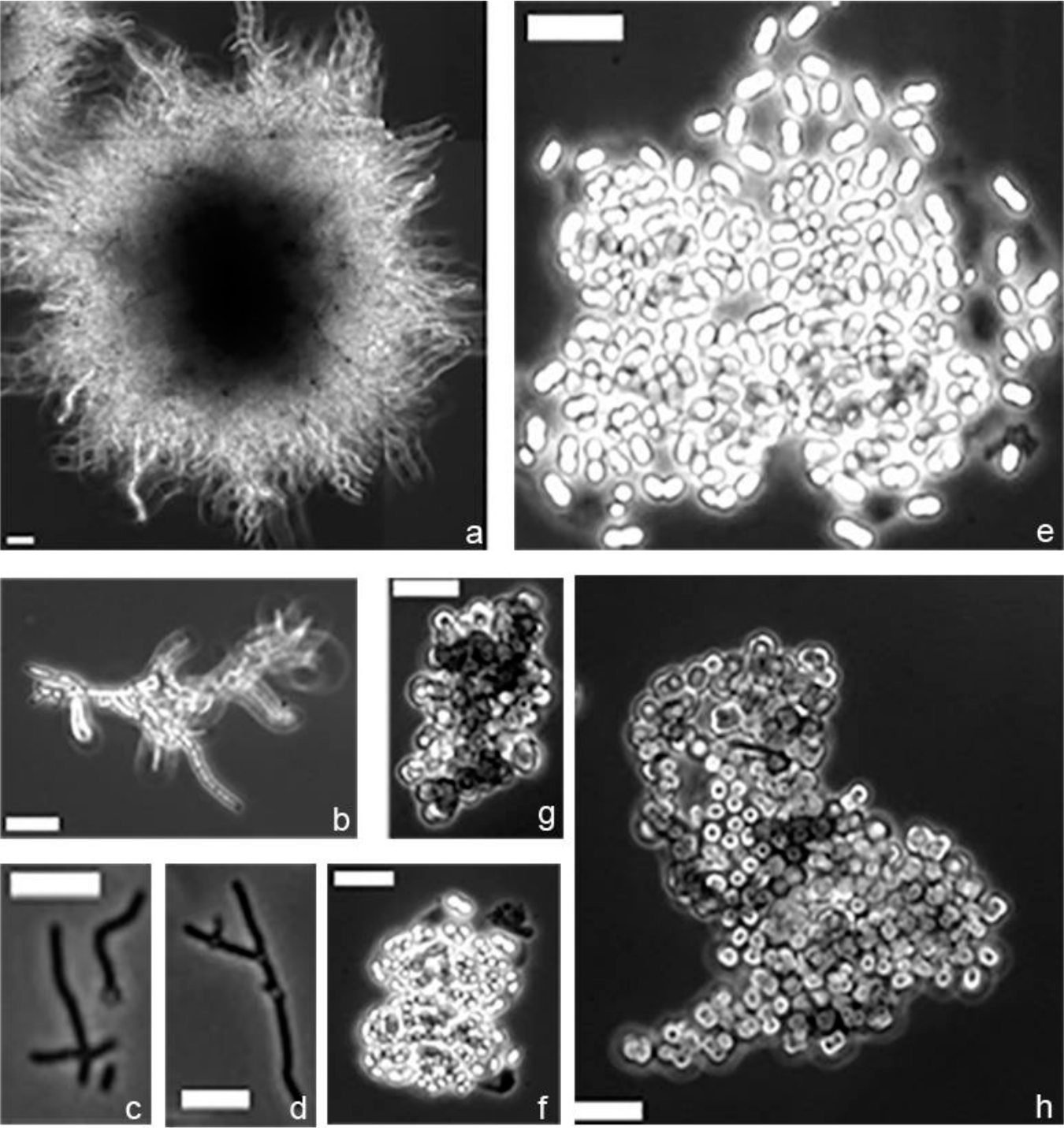
Application of SParticle to image analysis for other filamentous organisms. Mycelial fragments and pellets were identified in cultures of *Aspergillus niger* (a, b), *Synechococcus elongates* PCC 7942 (c, d), *Gloethece* sp. PCC6909 (e, f), and *Chroococcidiopsis* sp. PCC 6712 (g, h). Scale bar, 5 µm.

## DISCUSSION

*Streptomyces* species are exploited extensively as production hosts of natural products and enzymes. However, their complex morphology in submerged cultures such as during industrial fermentation significantly compromises their commercialization. Understanding the relationship between morphology and productivity of antibiotics and heterologous enzymes should, in part, identify how a strain may be morphologically tuned for optimal yield. Multiple methods have been applied to alter pellet morphology, based on changing environmental factors that alter growth or fragmentation (Mehmood 2011; Wucherpfennig 2011), on altering morphological genes such as those for surface polysaccharides (*cslA, glxA, matAB*; (Chaplin 2015; van Dissel 2015; Xu 2008)) or for cell division genes (*ssgA*; (Traag and van Wezel 2008; van Wezel 2006)). Many of these morphological alterations have been shown to influence and often improve yield and productivity. This stresses the need for morphological optimization of *Streptomyces* cultures, where the relation between small changes in morphology and product formation is monitored. The new imaging tool developed during this work allows for detailed and at the same time rapid assessment of particle sizes and shapes in a given culture, thus enabling scientists to study the relationship between morphology and productivity efficiently and in high detail.

WSI-based image analysis represents a quick, easy and inexpensive method for analysis of pellet shapes and heterogeneity in *Streptomyces* cultures. The only specialized equipment that is needed is an automated XY-stage for a wide-field microscope. If such equipment, which is often routinely present in many imaging setups, is not available, the method can be used to automate the analysis of single images, although in that care must be taken to prevent subjective image selection. Besides the ease of implementation, our method can also be adapted to include other microscopic methods, whereby qualitative pellet analysis may be based on for example fluorescence, such as for the study of mycelial aggregation (Zacchetti 2016). In addition, the tool may be adapted for the analysis of filamentous bacteria in activated sludge processes where current tools are only 72% accurate (Dias 2016).

Imaging mycelia formed after 24 h and 48 h of growth with different inoculums were well in accordance with previously described dynamics. Imaging of a single slide containing up to 500 pellets and mycelial fragments can be done in less than 10 min. While this does not compare to the high rate of flow cytometry-based methods (COPAS), the throughput is high enough to analyze population dynamics and has the added advantage of being retraceable because the original images can be reanalyzed. Also, a COPAS is expensive, and the low-cost WSI setup makes the method widely applicable, and at much higher throughput than manual microscopic methods. Additional flexibility is offered by the fact that additional filters can be generated using the Roi_Manager_Filter_Creator tool. This allows creating a storable and reusable Roi-Manager filter allowing the user to set their own filtering parameters. Thus, the program can be adapted to analyze basically anything that can be separated from the background of given sample image by using intensity-based thresholding. The combination of tools should be widely applicable to anyone interested in automated image analysis of whole slide imaging.

## Acknowledgements

We are grateful to Gerben Voshol for providing *Aspergillus niger* N402, *Gloeothece sp.* PCC 6909, *Chroococcidiopsis sp.* PCC 6712 and *Synechococcus elongatus* PCC 7942. The work was supported by grants from the Netherlands Organization for Scientific Research (NWO VICI grant 10379; VIDI grant 12957).

## List of abbreviations

AR: Aspect ratio
B&W: Black & white
Cfu: Colony forming units
cm: centimeter
COPAS: Complex Object Parametric Analyzer and Sorter
Gb: Gigabyte
h: hours
mL: milliliter
MS: Mannitol soy flour agar
nm: nanometer
RAM: Random access memory
ROI: Region of interest
Rpm: rounds per minute
TIFF: tag image file format
TSBS: Tryptic soy broth with sucrose
WSI: Whole slide imaging
µm: micrometer

## References

Barka EA, Vatsa P, Sanchez L, Gavaut-Vaillant N, Jacquard C, Meier-Kolthoff J, Klenk HP, Clément C, Oudouch Y, van Wezel GP (2016) Taxonomy, physiology, and natural products of the *Actinobacteria*. Microbiol Mol Biol Rev 80: 1–43.

Barry DJ, Williams GA (2011) Microscopic characterisation of filamentous microbes: towards fully automated morphological quantification through image analysis. J Microsc 244: 1–20.

Bentley SD, Chater KF, Cerdeno-Tarraga AM, Challis GL, Thomson NR, James KD, Harris DE, Quail MA, Kieser H, Harper D, Bateman A, Brown S, Chandra G, Chen CW, Collins M, Cronin A, Fraser A, Goble A, Hidalgo J, Hornsby T, Howarth S, Huang CH, Kieser T, Larke L, Murphy L, Oliver K, O’Neil S, Rabbinowitsch E, Rajandream MA, Rutherford K, Rutter S, Seeger K, Saunders D, Sharp S, Squares R, Squares S, Taylor K, Warren T, Wietzorrek A, Woodward J, Barrell BG, Parkhill J, Hopwood DA (2002) Complete genome sequence of the model actinomycete *Streptomyces coelicolor* A3(2). Nature 417: 141–147.

Bérdy J (2005) Bioactive microbial metabolites. J Antibiot (Tokyo) 58: 1–26.

Celler K, Picioreanu C, van Loosdrecht MC, van Wezel GP (2012) Structured morphological modeling as a framework for rational strain design of Streptomyces species. Antonie Van Leeuwenhoek 102: 409–423.

Chaplin AK, Petrus ML, Mangiameli G, Hough MA, Svistunenko DA, Nicholls P, Claessen D, Vijgenboom E, Worrall JA (2015) GlxA is a new structural member of the radical copper oxidase family and is required for glycan deposition at hyphal tips and morphogenesis of *Streptomyces lividans*. Biochem J 469: 433–444.

Chater KF, Losick R (1997) Mycelial life style of *Streptomyces coelicolor* A3(2) and its relatives. In: Shapiro JA, Dworkin M (eds) Bacteria as multicellular organisms. Oxford University Press, New York, pp 149–182.

Claessen D, Rozen DE, Kuipers OP, Sogaard-Andersen L, van Wezel GP (2014) Bacterial solutions to multicellularity: a tale of biofilms, filaments and fruiting bodies. Nat Rev Microbiol 12: 115–124.

Cooper LA, Kong J, Gutman DA, Dunn WD, Nalisnik M, Brat DJ (2015) Novel genotype-phenotype associations in human cancers enabled by advanced molecular platforms and computational analysis of whole slide images. Lab Invest 95: 366–376.

Cox PW, Thomas CR (1992) Classification and measurement of fungal pellets by automated image analysis. Biotechnol Bioeng 39: 945–952.

Cui YQ, Okkerse WJ, van der Lans RG, Luyben KC (1998) Modeling and measurements of fungal growth and morphology in submerged fermentations. Biotechnol Bioeng 60: 216–229.

Dias PA, Dunkel T, Fajado DA, Gallegos Ede L, Denecke M, Wiedemann P, Schneider FK, Suhr H (2016) Image processing for identification and quantification of filamentous bacteria in in situ acquired images. Biomed Eng Online 15: 64.

Girard G, Traag BA, Sangal V, Mascini N, Hoskisson PA, Goodfellow M, van Wezel GP (2013) A novel taxonomic marker that discriminates between morphologically complex actinomycetes. Open Biol 3: 130073.

Glazebrook MA, Doull JL, Stuttard C, Vining LC (1990) Sporulation of *Streptomyces venezuelae* in submerged cultures. J Gen Microbiol 136: 581–588.

Gray A, Wright A, Jackson P, Hale M, Treanor D (2015) Quantification of histochemical stains using whole slide imaging: development of a method and demonstration of its usefulness in laboratory quality control. J Clin Pathol 68: 192–199.

Hardy N, Moreaud M, Guillaume D, Augier F, Nienow A, Béal C, Ben Chaabane F (2017) Advanced digital image analysis method dedicated to the characterization of the morphology of filamentous fungus. J Microsc.

Hopwood DA (2007) Streptomyces in nature and medicine: the antibiotic makers. Oxford University Press, New York.

Jakimowicz D, van Wezel GP (2012) Cell division and DNA segregation in *Streptomyces:*how to build a septum in the middle of nowhere? Mol Microbiol.

Kendrick KE, Ensign JC (1983) Sporulation of *Streptomyces griseus* in submerged culture. J Bacteriol 155: 357–366.

Kieser T, Bibb MJ, Buttner MJ, Chater KF, Hopwood DA (2000) Practical Streptomyces genetics. John Innes Foundation, Norwich, U.K.

Kington JDI (2012) The Structure and Kinematics of the Nankai Trough Accretionary Prism, Japan. UNIVERSITY OF WISCONSIN-MADISON,

Mehmood N, Olmos E, Goergen JL, Blanchard F, Ullisch D, Klockner W, Buchs J, Delaunay S (2011) Oxygen supply controls the onset of pristinamycins production by Streptomyces pristinaespiralis in shaking flasks. Biotechnol Bioeng 108: 2151–2161.

Meyerhoff J, Tiller V, Bellgardt KH (1995) Two mathematical models for the development of a single microbial pellet. Biotechnol Bioeng 12: 305–313.

Obert M, Pfeifer P, Sernetz M (1990) Microbial growth patterns described by fractal geometry. J Bacteriol 172: 1180–1185.

Omura S (1992) Thom Award Lecture. Trends in the search for bioactive microbial metabolites. J Ind Microbiol 10: 135–156.

Packer HL, Thomas CR (1990) Morphological measurements on filamentous microorganisms by fully automatic image analysis. Biotechnol Bioeng 35: 870–881.

Papagianni M (2014) Characterization of Fungal Morphology using Digital Image AnalysisTechniques. Journal of Microbial & Biochemical Technology 6: 189–194

Petrus ML, van Veluw GJ, Wösten HA, Claessen D (2014) Sorting of Streptomyces cell pellets using a complex object parametric analyzer and sorter. J Vis Exp: e51178.

Pinto LS, Vieira LM, Pons MN, Fonseca MM, Menezes JC (2004) Morphology and viability analysis of Streptomyces clavuligerus in industrial cultivation systems. Bioprocess Biosyst Eng 26: 177–184.

Posch AE, Spadiut O, Herwig C (2012) A novel method for fast and statistically verified morphological characterization of filamentous fungi. Fungal Genet Biol 49: 499–510.

Reichl U, Buschulte TK, Gilles ED (1990) Study of the early growth and branching of Streptomyces tendae by means of an image processing system. J Microsc 158: 55–62.

Reichl U, King R, Gilles ED (1992) Characterization of pellet morphology during submerged growth of Streptomyces tendae by image analysis. Biotechnol Bioeng 39: 164–170.

RØnnest NP, Stocks SM, Lantz AE, Gernaey KV (2012) Comparison of laser diffraction and image analysis for measurement of Streptomyces coelicolor cell clumps and pellets. Biotechnol Lett 34: 1465–1473.

Ryoo D (1999) Fungal fractal morphology of pellet formation in Aspergillus niger. Biotechnology techniques 13: 33–36.

Stojmenovic M, Jevremovic A, Nayak A Fast iris detection via shape based circularity. In: Industrial Electronics and Applications (ICIEA), 2013 8th IEEE Conference on, 19-21 June 2013 2013. pp 747–752.

Tough AJ, Prosser JI (1996) Experimental verification of a mathematical model for pelleted growth of Streptomyces coelicolor A3(2) in submerged batch culture (vol 142, pg 639, 1996). Microbiol-Uk 142: 1332–1332.

Traag BA, van Wezel GP (2008) The SsgA-like proteins in actinomycetes: small proteins up to a big task. Antonie Van Leeuwenhoek 94: 85–97.

Treskatis SK, Orgeldinger V, Wolf H, Gilles ED (1997) Morphological characterization of filamentous microorganisms in submerged cultures by on-line digital image analysis and pattern recognition. Biotechnol Bioeng 53: 191–201.

Tresner HD, Hayes JA, Backus EJ (1967) Morphology of Submerged Growth of Streptomycetes as a Taxonomic Aid.I. Morphological Development of *Streptomyces Aureofaciens* in Agitated Liquid Media. Appl Microbiol 15: 1185-&.

van Dissel D, Claessen D, Roth M, van Wezel GP (2015) A novel locus for mycelial aggregation forms a gateway to improved *Streptomyces* cell factories. Microb Cell Fact: In press.

van Dissel D, Claessen D, Van Wezel GP (2014) Morphogenesis of *Streptomyces* in submerged cultures. Adv Appl Microbiol 89: 1–45.

van Wezel GP, Krabben P, Traag BA, Keijser BJ, Kerste R, Vijgenboom E, Heijnen JJ, Kraal B (2006) Unlocking *Streptomyces* spp. for use as sustainable industrial production platforms by morphological engineering. Appl Environ Microbiol 72: 5283–5288.

van Wezel GP, McKenzie NL, Nodwell JR (2009) Chapter 5. Applying the genetics of secondary metabolism in model actinomycetes to the discovery of new antibiotics. Methods Enzymol 458: 117–141.

Vecht-Lifshitz SE, Magdassi S, Braun S (1990) Pellet formation and cellular aggregation in Streptomyces tendae. Biotechnol Bioeng 35: 890–896.

Vinck A, Terlou M, Pestman WR, Martens EP, Ram AF, van den Hondel CA, Wosten HA (2005) Hyphal differentiation in the exploring mycelium of *Aspergillus niger*. Mol Microbiol 58: 693–699.

Voshol GP, Meyer V, van den Hondel CA (2015) GTP-binding protein Era: a novel gene target for biofuel production. BMC Biotechnol 15: 21.

Wardell JN, Stocks SM, Thomas CR, Bushell ME (2002) Decreasing the hyphal branching rate of *Saccharopolyspora erythraea* NRRL 2338 leads to increased resistance to breakage and increased antibiotic production. Biotechnol Bioeng 78: 141–146.

Webster JD, Dunstan RW (2014) Whole-slide imaging and automated image analysis: considerations and opportunities in the practice of pathology. Vet Pathol 51: 211–223.

Willemse J, Ruban-Osmialowska B, Widdick D, Celler K, Hutchings MI, van Wezel GP, Palmer T (2012) Dynamic localization of Tat protein transport machinery components in *Streptomyces coelicolor*. J Bacteriol 194: 6272–6281.

Willemse J, van Wezel GP (2009) Imaging of Streptomyces coelicolor A3(2) with reduced autofluorescence reveals a novel stage of FtsZ localization. PLoS One 4: e4242.

Wucherpfennig T, Hestler T, Krull R (2011) Morphology engineering‒‒osmolality and its effect on Aspergillus niger morphology and productivity. Microb Cell Fact 10: 58.

Wucherpfennig T, Lakowitz A, Krull R (2013) Comprehension of viscous morphology- evaluation of fractal and conventional parameters for rheological characterization of Aspergillus niger culture broth. J Biotechnol 163: 124–132.

Xu H, Chater KF, Deng Z, Tao M (2008) A cellulose synthase-like protein involved in hyphal tip growth and morphological differentiation in Streptomyces. J Bacteriol 190: 4971–4978.

Zacchetti B, Willemse J, Recter B, van Dissel D, van Wezel GP, Wosten HA, Claessen D (2016) Aggregation of germlings is a major contributing factor towards mycelial heterogeneity of *Streptomyces*. Sci Rep 6: 27045.

